# A novel nanoluciferase transgenic reporter to measure proteinuria in zebrafish

**DOI:** 10.1101/2021.07.19.452884

**Authors:** Richard W. Naylor, Emmanuel Lemarie, Anthony Jackson-Crawford, J. Bernard Davenport, Aleksandr Mironov, Martin Lowe, Rachel Lennon

## Abstract

The zebrafish is an important animal system for modelling human diseases. This includes kidney dysfunction as the embryonic kidney (pronephros) shares considerable molecular and morphological homology with the human nephron. Zebrafish also have a high fecundity, with females capable of laying 200-300 eggs per week, thereby facilitating chemical and mutation screening. A key clinical indicator of kidney disease is proteinuria, but a high-throughput readout of proteinuria in the zebrafish is lacking. Coupling the advantages of the zebrafish system with a tool to measure proteinuria will advance the scope for testing the efficacy of drugs to treat kidney diseases. Here, we generated a stable transgenic zebrafish line using the *l-fabp10* liver-specific promoter to over-express a nanoluciferase molecule fused with the D3 domain of Receptor-Associated-Protein (RAP) to create NL-D3. In the healthy state, NL-D3 is excreted, but when embryos were treated with chemicals that affected either proximal tubular reabsorption (cisplatin, gentamicin) or glomerular filtration (angiotensin II, Hanks Balanced Salt Solution, Bovine Serum Albumin), NL-D3 presence in the urine increased. Similarly, depletion of several gene products associated with kidney disease (*nphs1, nphs2, lrp2a, ocrl, col4a3, col4a4*, and *col4a5*) also induced NL-D3 proteinuria. Furthermore, we found that treating *col4a4* depleted zebrafish larvae (a model of Alport syndrome) with captopril reduced proteinuria. Our findings confirm the use of the *NL-D3* transgenic zebrafish as a robust and quantifiable proteinuria reporter. Given the feasibility of high-throughput assays in zebrafish, this novel reporter will permit screening for drugs that ameliorate proteinuria and thereby prioritise candidates for further translational studies.

**Significance Statement:** The zebrafish has become an important system for modelling kidney disease. However, proteinuria, an important clinical indicator of kidney dysfunction, is not easily detected in zebrafish. Here, we describe a transgenic line that uses a nanoluciferase reporter to enable detection of proteinuria in multiple models of glomerular and proximal tubular kidney disease in the zebrafish. In this system proteinuria can be accurately measured in a high-throughput manner and will enable the screening of drugs that affect glomerular filtration or protein re-uptake in the proximal tubule.

## Introduction

Proteinuria is a key clinical indicator of kidney disease (National Kidney Foundation, 2002). Using urine dip sticks, the presence of protein in the urine can be determined in seconds. Further evaluation of the size of protein in the urine indicates the site of functional deficit along the nephron; the presence of large proteins in the urine suggests dysfunction in the glomerular filter, whereas the presence of low molecular weight proteins indicates proximal tubule dysfunction (Butt et al., 2020; Lawrence et al., 2017). Whilst the use of proteinuria as an indicator of kidney function is admissible in humans, given the ease of urine acquisition and the relatively large volume of urine produced, measuring proteinuria (and therefore kidney function) in lab-based animal research is more difficult. This is particularly problematic in the zebrafish, which is a popular model organism for the study of kidney development and disease (Morales and Wingert, 2017; Outtandy et al., 2019; Poureetezadi and Wingert, 2016). A quantitative reporter of zebrafish proteinuria would therefore improve the toolkit available to kidney researchers.

Embryonic zebrafish develop a functioning pronephric kidney that has molecular and structural analogy to a human nephron (Drummond et al., 1998). The pronephros consists of a single midline fused glomerulus attached to bilateral tubules that are segmented along their anterior-posterior axis into distinct proximal and distal domains (Drummond et al., 1998; Wingert et al., 2007). The posterior component of the pronephros is a short duct-like region that plumbs into the cloacal opening to allow excretion. Zebrafish embryos do not have a bladder and so constantly excrete into the external environment. The lack of a bladder and the small volume of excreted urine makes it difficult to assay proteinuria in zebrafish in a reliable and quantitative manner. Until recently, the best method of determining kidney function in zebrafish embryos was the use of fluorescent dextrans of varying molecular weights (Christou-Savina et al., 2015). As with the human glomerulus, the size selectivity limit of the zebrafish glomerulus is ~70 kDa. Therefore, injected fluorescent dextrans of a larger size (for example 100 kDa or 500 kDa) should not pass through a healthy zebrafish glomerular capillary wall into the urinary space. The presence of these proteins in the proximal tubules (detected by fluorescence microscopy) indicates glomerular dysfunction. Similarly, fluorescent dextrans smaller than ~70 kDa should traverse the glomerular filtration barrier and be detected in endosomal structures within the proximal tubular epithelia. If these smaller dextrans are not observed in the proximal tubular epithelium, then this indicates proximal tubule dysfunction. Fluorescent dextran is not specific to the cellular machinery of protein endocytosis, it is passively internalised in the fluid bulk. As such, these methods are useful, but they are not quantitative, do not specifically identify megalin-mediated endocytosis defects in the proximal tubule, and cannot be used in a high-throughput manner.

The zebrafish is a valuable animal model system because it has high fecundity (200-300 eggs laid per week per female), is highly transparent, and development occurs *ex utero*. These characteristics along with its amenability to genetic manipulation, make it an ideal vertebrate system with which to perform high-throughput drug or mutation screens on genetically similar embryos (Gehrig et al., 2018; Parng et al., 2002). A precise and efficient method of quantifying proteinuria in the zebrafish would be a powerful tool for investigating the efficacy of new drugs or determining the potential nephrotoxicity of specific reagents. To this end, a zebrafish transgenic reporter (*l-fabp10:½vdbp-mCherry*) that enabled embryo medium to be collected and assayed for low molecular weight proteins was recently developed (Chen et al., 2020). This powerful system permits quantification of proteinuria as a readout of dysfunction of the lrp2/megalin endocytosis pathway in the proximal tubule. It reports proximal tubular dysfunction following downstream processing with immunofluorescence imaging or ELISA assays to quantify protein uptake in the proximal tubules and proteinuria. This genetic tool highlights the potential of transgenics to develop reporters of proteinuria in the zebrafish.

Here, we describe a novel nanoluciferase based reporter of proteinuria in zebrafish. Nanoluciferase is an excellent candidate marker due to its small size and 100-fold increased brightness relative to other luciferase enzymes. The level of excreted protein in the embryo medium is detected by luminescence of nanoluciferase following simple addition of the substrate luciferin. The intensity of luminescence signal can be used to measure the amount of nanoluciferase protein excreted by the zebrafish pronephros. Furthermore, this new system measures perturbations in both proximal tubular and glomerular function and its simple and easy application makes it amenable to use in a high-throughput manner.

## Methods

### Zebrafish husbandry

Zebrafish were maintained and staged according to established protocols (Kimmel et al., 1995) and in accordance with the project licenses of Martin Lowe (70/9091) and Rachel Lennon (P1AE9A736) under the current guidelines of the UK Animals Act 1986. Embryos were collected from group-wise matings of wild-type AB Notts or *y-crystallin:mcherry/l-fabp10:NL-D3* fish.

### Molecular Cloning

The cDNA for rat RAP was a gift from Prof. Thomas Willnow (Max Delbruck Centre for Molecular Medicine, Berlin). The nanoluciferase (NL) cDNA was from Promega, with an N-terminal fusion of the IL-6 signal sequence (Hall et al., 2012). The D3 fragment of RAP (amino acid numbers 206-319, lacking the carboxy-terminal HNEL ER retention sequence) was cloned downstream of the NL cDNA prior to the stop codon to create a C-terminal fusion. NL-D3 was cloned by PCR into the pTrcHis bacterial expression vector for preparation of recombinant His-tagged protein in *E. coli*. For generation of zebrafish transgenics, the Gateway cloning method was used (Kwan et al., 2007). NL-D3 cDNA was cloned into the Gateway p3 vector. The p1 vector contained the γ-Crystallin promoter driving mCherry expression (Offield et al., 2000) and p2 vector contained the liver specific *lfabp10* promoter (Her et al., 2003). The p1-p3 plasmids were combined with the p4-I-SceI SAR-CH4 Tol2 destination vector (Love et al., 2011) to make the final transgenesis construct.

### Zebrafish transgenesis

5 μl of 80 ng/μl DNA of the transgenesis construct were mixed with 5 μl of 50 ng/μl *tol2* transposase mRNA and 1 μl of 10X phenol red. All solutions were prepared fresh immediately before injection. Each single cell stage embryo was oriented to allow access into the cell and 1 nl of this solution was injected directly into the cell using a microinjector. Successful transgenesis was confirmed in F_0_ embryos by selecting for mCherry expression in the lens at day 5, and founder fish were then grown to adulthood.

### Preparation of recombinant NL-D3

His-tagged NL-D3 was expressed in *E. coli* BL21 (De3) Codon Plus cells by induction with 1 mM IPTG for 16 hours at 18°C. The protein was purified using Ni-NTA agarose using standard methods, snap frozen in liquid nitrogen, and stored at −80°C until use.

### Cell culture

Parental HEK293 EBNA cells or HEK293 EBNA cells stably expressing megalin mini-receptor (MmR4) were kindly provided by Prof. Thomas Willnow (Max Delbruck Centre for Molecular Medicine, Berlin). The cells were maintained at 37°C in 5% CO_2_ in DMEM containing 10% fetal bovine serum, penicillin (100 U/ml), streptomycin (0.1 mg/ml), and 0.35 mg/ml G418 without (parental) or with (MmR4 cells) an additional 1 μg/ml puromycin. For NL-D3 uptake, cells were washed with PBS, then serum-free DMEM before preincubating in serum-free DMEM for 30 min at 37°C. Cells were then incubated in serum-free DMEM containing NL-D3 at 1 μg/ml for 30 min at 37°C, washed 3x in PBS, and then lysed in Nano-Glo luciferase buffer (Promega) for assessment of luciferase activity.

### Drug treatments

Drug treatments were performed on zebrafish embryos either by injection into the common cardinal vein (HBSS (Thermofisher #14170070, neat), BSA (ThermoFisher #B14, 10%)) or by incubation in the E3 embryo medium (captopril (Bio-Techne #4455/50, 10 μM), angiotensin (Eurogentec #AS20634, 0.5-5 μM)). For injection, embryos were grown to desired stage (3 dpf) and then anaesthetised in buffered tricaine methanesulfonate (MS-222) at 164 mg/L. Embryos were then placed in an injection chamber containing MS-222 and injected into the common cardinal vein with a microinjection needle. Embryos were then transferred to fresh E3 embryo medium to recover and grown to desired stage. For incubations, embryos were transferred with a transfer pipette to fresh E3 embryo medium containing the drug at the desired concentration. Control embryos were treated with the equivalent concentration of vehicle substance (DMSO).

For gentamicin and cisplatin treatments, previous methods described by Chen et al, (2020) were used. In brief, gentamicin (Sigma, #G1264) was re-suspended in a 0.9% NaCl solution to a concentration of 10 ng/μl. This was diluted to 6 ng/μl for injection, which was performed at 48 hpf into the common cardinal vein (either 0.5 nl or 1 nl injection volume). For cisplatin treatments, the drug was re-suspended directly into E3 embryo medium containining 0.01% methylene blue. Concentrations of 0.5 mM and 1.5 mM were exposed to embryos at 48 hpf for a period of four hours. Embryos were then washed three times in E3 medium and left to grow to the desired stage for proteinuria analysis.

### Morpholino oligonucleotide treatments

For *lrp2a* and *ocrl* morpholino experiments, both morpholinos were diluted to a final concentration of 0.24 mM. For control experiments, 0.24 mM of a standard control morpholino oligonucleotide was used. For *lrp2a* and *ocrl* knockdowns, a *p53* morpholino was also co-injected at the same concentration (0.24 mM). For *nphs1* and *nphs2* morpholino experiments, final concentrations of 0.15 mM (*nphs1*^MOex25^) and 0.25 mM (*nphs2*^MOex3^) were used. Injection droplet sizes were 5 nl for all morpholino experiments, and if defrosting a morpholino, the oligonucleotide was heated at 65°C for 10 minutes prior to adding to the injection mixture. All microinjections were into the cell compartment of one-cell stage embryos. Morpholino sequences are shown below;

*standard control* – 5’ CCTCTTACCTCAGTTACAATTTATA 3’
*p53* – 5’ GCGCCATTGCTTTGCAAGAATTG 3’
*lrp2a* – 5’ AATCAGTGCTTGTGGTTTACCTGGG 3’
*ocrl* – 5’ AATCCCAAATGAAGGTTCCATCATG 3’
*nphs1* – 5’ TGCACCAACACGACTCACCTCTGCTC 3’
*nphs2* – 5’ TGTAGTCACTTTTGCAGACCTGGGCT 3’

### CRISPR-Cas9 knockdowns

To genetically knockdown expression of gene products, we adopted the CRISPR-Cas9 approach described by Wu et al., (2018). gRNAs were order from Merck and resuspended in nuclease-free water to a concentration of 20 μM. For the injection mix, 4 μM of each gRNA was combined with Cas9 (NEB #M0646) and Cas9 buffer. This injection mixture was then injected into the cytoplasm of 1-cell stage NL-D3 transgenic embryos. Below are the gRNAs used in this work;

col4a3 gRNA1 - GGTCGGAGCATGGGAATACC
col4a3 gRNA1 - CGTTGGCCATCACGACCTTT
col4a3 gRNA1 - GAAATAGCATTTGGCTGCCC
col4a3 gRNA1 - TGTTGAATGAGTCACCTGGT

col4a4 gRNA1 - GATCCAGGACTATCTTTACC
col4a4 gRNA2 - CTTGAAGCCCTTTAGGGCCA
col4a4 gRNA3 - GGATACCCCGGTGTTCCCGG
col4a4 gRNA4 - TGGATCCCCCATTAACCCCA

col4a5 gRNA1 - GACGGACCTCAAGGGTCGAT
col4a5 gRNA2 - GTCCAGGTATTCCCTAAGGT
col4a5 gRNA3 - GGTCCTGGCAAACCAATACC
col4a5 gRNA4 - GGAAATCCAGCACTACCGGC

control gRNA1 - CAGGCAAAGAATCCCTGCC
control gRNA2 - TACAGTGGACCTCGGTGTC
control gRNA3 - CTTCATACAATAGACGATG
control gRNA4 - TCGTTTTGCAGTAGGATCG

### Fluorescent re-uptake assays

β-Lactoglobulin from bovine milk (Sigma, L3908) was resuspended in 1X PBS and labelled with Cy3 mono-reactive dye (Amersham, PA23001) according to the manufacturer’s instructions. 5 dpf embryos were anesthetised in E3 and MS222 (0.4 mg/ml) solution and then transferred to a 1% agarose injection dish. Larvae were injected in the common cardinal vein with 1 nl of Cy3-β-Lactoglobulin (5 mg/ml) and/or Alexa fluor™ 488 10 kDa dextran (D22910, Thermo Fisher, at 2 mg/ml) diluted in sterile H_2_O and successful injections were screened for on a Leica Mz10F fluorescent stereomicroscope. Uptake in the proximal tubule was visualised <1 hour post injection.

### Standard curve generation

Fifty microlitres of serially diluted recombinant NL-D3 protein in PBS (0.5 ng/ml to 0.0005 ng/ml) were placed in the wells of a 96-well plate. The Nano-Glo^®^ luciferase assay system (Promega #N1110) was used, with 50 μl of this solution added to each well containing recombinant NL-D3, or the buffer only blank. The plate was then placed in a Flexstation^®^ 3 Multi-mode microplate reader and luminescence was measured by reading each well for 1 second. Data was collected in relative luminescence units (RLU) using SoftMax ^®^ Pro software. Using RLU values of each serial dilution minus the blank, a standard curve of known concentrations of recombinant NL-D3 relative to RLU was established.

### Zebrafish proteinuria reporter assay

The pipeline of the assay for proteinuria in *NL-D3* zebrafish is shown in the schematics accompanying Figures 2B and 6B. Embryos were grown to 4 dpf, then three embryos per well were placed in one well of a 96-well dish (Figure 2B). E3 embryo medium was removed and replaced with 200 μl fresh E3. 24 hours later 50 μl of E3 medium was removed from each well and placed in the corresponding well of a fresh opaque 96-well plate. 50 μl of substrate from the NanoGlo^®^ Luciferase Assay System (Promega #N1110) was then added to each well. Plates were then briefly spun down at 700 rpm for 1 minute and then immediately assayed for luminescence on a Flexstation 3 multi-mode microplate reader. Softmax Pro 5.4 software (Molecular Devices) was used to detect luminescence. Endpoint analysis was selected with a 1000 millisecond integration of luminescence (RLUs). Opaque 96-well option was selected in ‘Assay Plate’, as well as Costar 96 opaque 3mL in ‘Compound Source’.

### RT-PCR

For RT-PCR, cDNA was diluted to 1 ng/μl and mixed with 7.8 μM of primers and added in equal volume to 2X *Power* SYBR™ Green PCR Master Mix (Thermofisher #4367659). The primers used are described below;

actb F - TCACCACCACAGCCGAAAG
actb R - AGAGGCAGCGGTTCCCAT
col4a3 F - AACTTGTCGCTACGCCTCTC
col4a3 R - ATTGCCTCGCATACGGAACA
col4a4 F - CTGGCTTTAAGGGACCTCCG
col4a4 R - AAGCAGACTGTAGCCGTTCC
col4a5 F - CCCAGGTCTTCCAGGACAGG
col4a5 R – CGATGACTTCCCCAGGTTTTGC

The PCR reaction was run on a Bio-Rad CFX96 Touch Real-Time PCR machine. All analyses of the data used the ΔΔCt method of quantification. RT-PCRs were run in triplicate with -RT and -cDNA controls.

### TIDE analysis

Embryos injected with gRNAs were grown to 24 hpf, dechorionated, then ten randomly selected embryos were placed in a 1.5 ml capped tube. All excess E3 embryo medium was removed and replaced with 50 μl of 50 μM NaOH. Embryos were then placed at 95°C in a heating block for 15 minutes. After a brief spin, 5 μl of Tris-HCl pH 8 was added to neutralize the samples. The genomic DNA preps were then homogenized by pipetting before 1 μl was used in a PCR to amplify the region around the gRNA cut sight using the primers shown in the primer list. The PCR product was purified, ran on a gel to check size, and then diluted to 50 ng/μl before being sequenced. Wild-type *control*^gRNA^-injected control ab1 files were used to compare crispants according to the TIDE guidelines (Brinkman et al., 2014). TIDE analysis was performed online at https://tide.nki.nl/

*col4a3* TIDE F – tgccaggtgttcagggaaat
*col4a3* TIDE R – aggctcctttggtcctgatg
*col4a4* TIDE F – tttctcaccccactcctcac
*col4a4* TIDE R – gatggcggtacacatggatg
*col4a5* TIDE F – atggtttgttctgttgactttca
*col4a5* TIDE R – gaatcttatgccggcccatg

### In situ hybridisation

Whole-mount in situ hybridisation on zebrafish embryos was performed as previously described (Thisse and Thisse, 2008). Digoxigenin-labelled anti-sense riboprobes were made using T3 RNA polymerase transcription kits (Roche Diagnostics). Probe templates for zebrafish *col4a3, col4a4* and *col4a5* were generated by PCR amplification from cDNA taken from 4 dpf zebrafish embryos. The primers used were as follows:

F *col4a3* 5’-AAAGGGGCTTGTGATTGCAG-3’,
R *col4a3* 5’-GGATCCAATTAACCCTCACTAAAGGGATCTCCAGCTCTGCCTTGTT-3’;
F *col4a4* 5’-CAGAGGCTTAACAGGTCCCA-3’,
R *col4a4* 5’-GGATCCAATTAACCCTCACTAAAGGGTCCACAATCCCCAGGTTCTC-3’.
F *col4a5* 5’-GCCTATTGTCTTGAAGGGCG-3’
R *col4a5* 5’-GGATCCAATTAACCCTCACTAAAGGGACGGGAAGCGAAGTTACAGA-3’

A T3 anchor sequence (GGATCCAATTAACCCTCACTAAAGGG) on the 5’ end of the reverse primer was used to enable T3-mediated RNA synthesis from the purified PCR product. Following colorimetric assay, the embryos were treated with 100% Methanol for 10 minutes with rocking to clear background labelling and were then fixed in 4% PFA for downstream imaging. Whole embryos were imaged on a Leica M205 FA upright stereofluorescence microscope, and transverse sections were imaged on an Olympus BX63 snapshot slide scanner microscope.

### Cryosectioning and immunofluorescence

4 dpf zebrafish embryos were fixed in Dent’s fixative (80% Methanol, 20% DMSO) or 4% PFA (for podocin staining) overnight at 4°C. These embryos were then washed three times in 1X PBS before being placed in cryomolds. All excess PBS was removed with a pipettor and finally with a Kimwipe. The cryomold chamber was then filled with OCT compound. The embryos were then oriented in one direction in the OCT. A beaker of isopentane prechilled (for at least 30 minutes) in a polystyrene box filled with dry ice was prepared. The cryomold was carefully submerged into the isopentane to flash freeze the sample. Samples were then stored at −80°C. Cryosectioning was performed in a Leica CM1950 cryostat, with 10 μm sections collected on coated glass slides. These collected sections were left to dry overnight at 4°C and then were processed for immunostaining. Slides were washed four times in PBS containing 1% Triton X100 (PBSTr). They were then washed once with distilled water before being permeabilized in acetone for 8 minutes at −20°C. Permeabilized sections were then washed in distilled water and four more times with PBSTr. The samples were then placed in blocking solution (5% BSA, 3% donkey serum in PBSTr) for at least one hour. The blocking solution was removed and replaced with the primary antibody in blocking solution. The primary antibodies used were NPHS2 (Abcam #ab50339; 1:250) and pan collagen IV (Abcam #ab6586; 1:250). Importantly, embryos fixed in PFA were used for the podocin staining and embryos fixed in Dent’s fixative were used for the pan collagen IV staining. The primary antibodies were left on overnight at 4°C and then washed at least 6x 1hr in PBSTr before being placed in secondary antibody Alexa Fluor 488 (1:500 in PBSTr) and left overnight again at 4°C. The slides were then washed five times in PBS containing 0.05% Tween before being mounted with a coverslip using Prolong Diamond antifade mountant (ThermoFisher #P10144). Images were collected on a Leica TCS SP8 AOBS inverted confocal using a 60X Plan Fluotar objective. The confocal settings were as follows, pinhole 1 airy unit, scan speed 1000Hz unidirectional, format 1024 × 1024. Images were collected using the white light laser with 488nm (10%) laser lines.

### Transmission electron microscopy

Samples were prepared according to protocols described previously (Randles et al., 2016). Images were taken on T12 Biotwin transmission electron microscope. Distances were measured in Fiji/ImageJ using a grid method, 133 measurements were taken and normalized to the length of the glomerular basement membrane.

### Statistical analysis

The mean ±SD was calculated using GraphPad Prism version 9 for Windows, GraphPad Software, San Diego, California USA, www.graphpad.com. Statistical significance scores were measured in GraphPad Prism version 9 using unpaired parametric Student’s t test. P values in 95% confidence limits were characterised as significant, R squared scores from this statistical analysis was also noted to determine the size of the difference between the two compared datasets.

## Results

### *Recombinant NL-D3 can be endocytosed* in vitro

We aimed to develop a transgenic reporter to study endocytosis in the zebrafish pronephric proximal tubule. An important component of endocytic pathways in the proximal tubule is the giant 600 kDa megalin receptor, which has multiple ligands (Christensen and Birn, 2002). Megalin function and processing is dependent on Receptor-Associated-Protein (RAP) (Birn et al., 2000), which contains three D-domains that bind with varying affinity to megalin (Medved et al., 1999) (**Figure 1A**). For our reporter protein, we fused rat RAP to nanoluciferase (NL). A full-length RAP-NL fusion yields a protein that is 58 kDa in size (**Figure 1A**). We hypothesized that limiting the size of the reporter protein would aid its transit through the glomerular capillary wall and so we fused the 14 kDa D3 domain of RAP to the C-terminus of NL to create NL-D3, which has a predicted size of 35.5 kDa (**Figure 1A**). We first wanted to confirm that the NL-D3 fusion protein could report megalin-dependent endocytosis. For this purpose, we expressed recombinant NL-D3 in *E. coli* and used affinity isolation to purify the protein (**Figure 1B**). The recombinant NL-D3 was then used in uptake experiments on HEK293 cells stably expressing a truncated version of megalin (containing the cytoplasmic tail, transmembrane domain and the fourth of four large extracellular repeats, named MmR4). Uptake was measured by lysing the cells and measuring luminescence and this showed that the HEK293-MmR4 cells incubated with recombinant NL-D3 had ~10-fold higher levels of uptake compared to control HEK293 cells incubated with recombinant NL-D3 (**Figure 1C**). Neither control HEK293 cells nor HEK293-MmR4 cells displayed luminescence after incubation with NL (**Figure 1C**). These *in vitro* results indicate that NL-D3 is bound by megalin and readily endocytosed.

**Figure 1:**
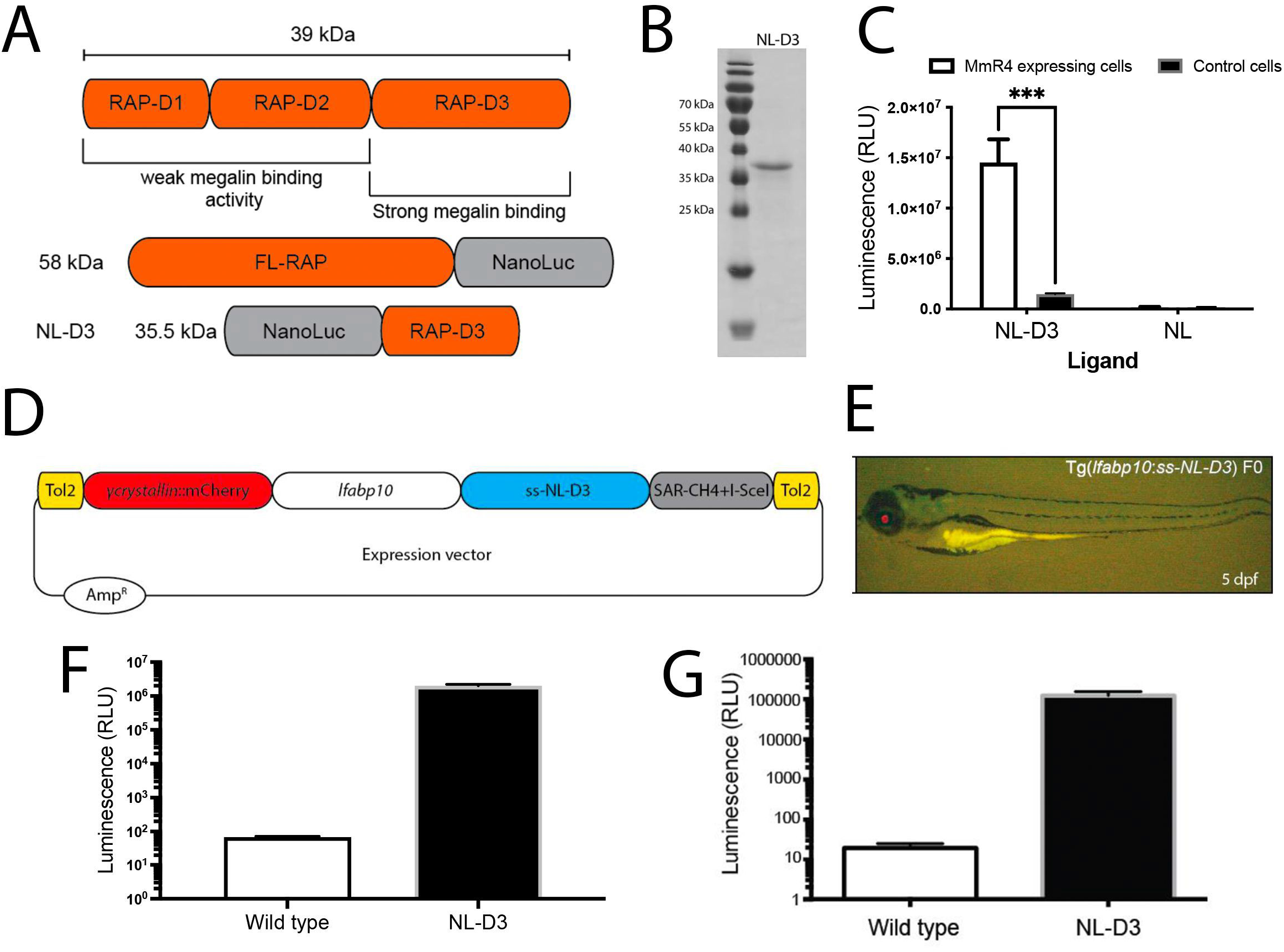
NL-D3 is uptaken by the megalin endocytosis pathway. A) Top schematic showing binding affinity to megalin of the three D-domains of the Receptor Associated Protein (RAP). Bottom schematics show the size and orientation of full-length RAP bound to the N-terminus of Nano-Luc and the RAP D3 domain bound to the C-terminus of Nano-Luc (NL-D3, which was used in this work). B) SDS-PAGE gel showing the recombinantly expressed and purified NL-D3 protein alongside molecular weight markers. C) Graph showing the internalisation of NL-D3 or untagged NL into MmR4 mini-megalin expressing cells and control non-megalin expressing cells. D) Schematic for the *y-crystallin:mcherry/l-fabp10:NL-D3* Tol2 vector used to generate transgenic zebrafish. E) Panel shows lateral view of a 5 dpf *NL-D3* transgenic zebrafish embryo under excitation for red fluorescence to highlight the mCh expression in the lens. F) Histogram showing relative luminescence units (RLU) in whole embryo lysates of wild-type and *NL-D3* 5 dpf embryos. G) Logarithmic histogram showing the RLU of 1 μl blood from adult wild-type and *NL-D3* zebrafish.

### *Development of* y-crystallin:mcherry/l-fabp10:NL-D3 *transgenic zebrafish*

To develop a viable *in vivo* reporter of proximal tubule endocytosis we turned to the zebrafish model. We used the versatile Tol2 gene transfer system (Kawakami, 2005) and Gateway™ cloning technology to generate transgenic zebrafish that express NL-D3 under the control of the liver-specific *l-fabp10* promoter (**Figure 1D**). To enable selection, the transposable element dually expressed mCherry under the control of the *y-crystallin* promoter, which allows transgenic embryos to be screened by assaying for red fluorescence in the lens (**Figure 1E**). Whole embryo lysates from stably transgenic 5 dpf *y-crystallin:mcherry/l-fabp10:NL-D3* zebrafish (hereon in termed *NL-D3*) had ~100,000-fold more luminescence than wild-type zebrafish embryos of the same developmental stage (**Figure 1F**). A similar increase in luminescence was observed in blood isolated from adult-stage *NL-D3* reporter fish (**Figure 1G**). We also measured the amount of NL-D3 in adult organ tissues, with the brain having relatively low amounts of NL-D3 compared to the kidney, which in turn had ~10-fold less NL-D3 than the liver (**Figure S1**). These results indicate liver-specific expression of NL-D3 in transgenic zebrafish results in the robust release of this reporter into the blood vasculature where it can then be filtered by the glomerulus.

### *Proteinuria is detected in* NL-D3 *embryos with proximal tubule and glomerular dysfunction*

To test if NL-D3 passes through the glomerular filter and is re-absorbed by the megalin endocytic pathway in the proximal tubule of the pronephros, we knocked down *lrp2a* (the gene that encodes for megalin) using a previously described morpholino oligonucleotide (Anzenberger et al., 2006). Depletion of *lrp2a* was sufficient to severely reduce uptake of a 10 kDa fluorescent dextran in proximal tubules (**Figure 2A**), confirming these morphant embryos have perturbed proximal tubule reabsorption. For the proteinuria assay, three morphant embryos at 4 days post fertilization (dpf) were placed in one-well of a 96-well plate (**Figure 2B**), the embryo medium was replaced, and embryos were cultured for an additional 24 hours before NL-D3 content in the embryo medium was measured on a luminometer. A standard curve was generated using recombinant NL-D3 to permit quantifiable conversion of relative luminescence units (RLUs) obtained from the luminometer to NL-D3 amount (ng/ml, **Figure S2**). Depletion of *lrp2a* caused a ~29-fold increase in NL-D3 present in the embryo medium (**Figure 2C**). Another gene product involved in proximal tubule endocytosis is *OCRL*, and pathogenic variants cause Lowe syndrome, which is associated with low molecular weight proteinuria (Bockenhauer et al., 2008; De Matteis et al., 2017). Morpholino knockdown of the zebrafish ortholog *ocrl* using a previously validated morpholino (Oltrabella et al., 2015; Ramirez et al., 2012) caused a reduction in 10 kDa fluorescent dextran uptake in the proximal tubule (**Figure 2A**), and induced proteinuria (~10-fold increase, **Figure 2C**). We also tested the effect of nephrotoxins that target proximal tubules and induce proteinuria. Gentamicin and cisplatin are two drugs that are well established nephrotoxins that cause renal damage primarily by proximal tubule cell death and they have previously been used to ablate the zebrafish proximal tubule epithelium (Hentschel et al., 2005). Analysis of NL-D3 secretion into embryo medium showed that 0.5 nl of 6 ng/μl gentamicin increased proteinuria by ~13-fold compared to vehicle controls (**Figure 2D**). Treating embryos with double the amount of gentamicin further increased the proteinuria to ~534-fold over controls (**Figure 2D**), suggesting a dose response is detectable in the *NL-D3* reporter. A similar dose-response was obtained with cisplatin, a low (0.5 mM) dose causing a ~1.5-fold increase in proteinuria, and a high (1.5 mM) dose causing a ~6.7-fold increase (**Figure 2E**). Taken together, these results confirm the *NL-D3* transgenic fish as a viable tool for detecting proteinuria due to proximal tubular dysfunction after genetic perturbation or pharmacological treatment.

**Figure 2:**
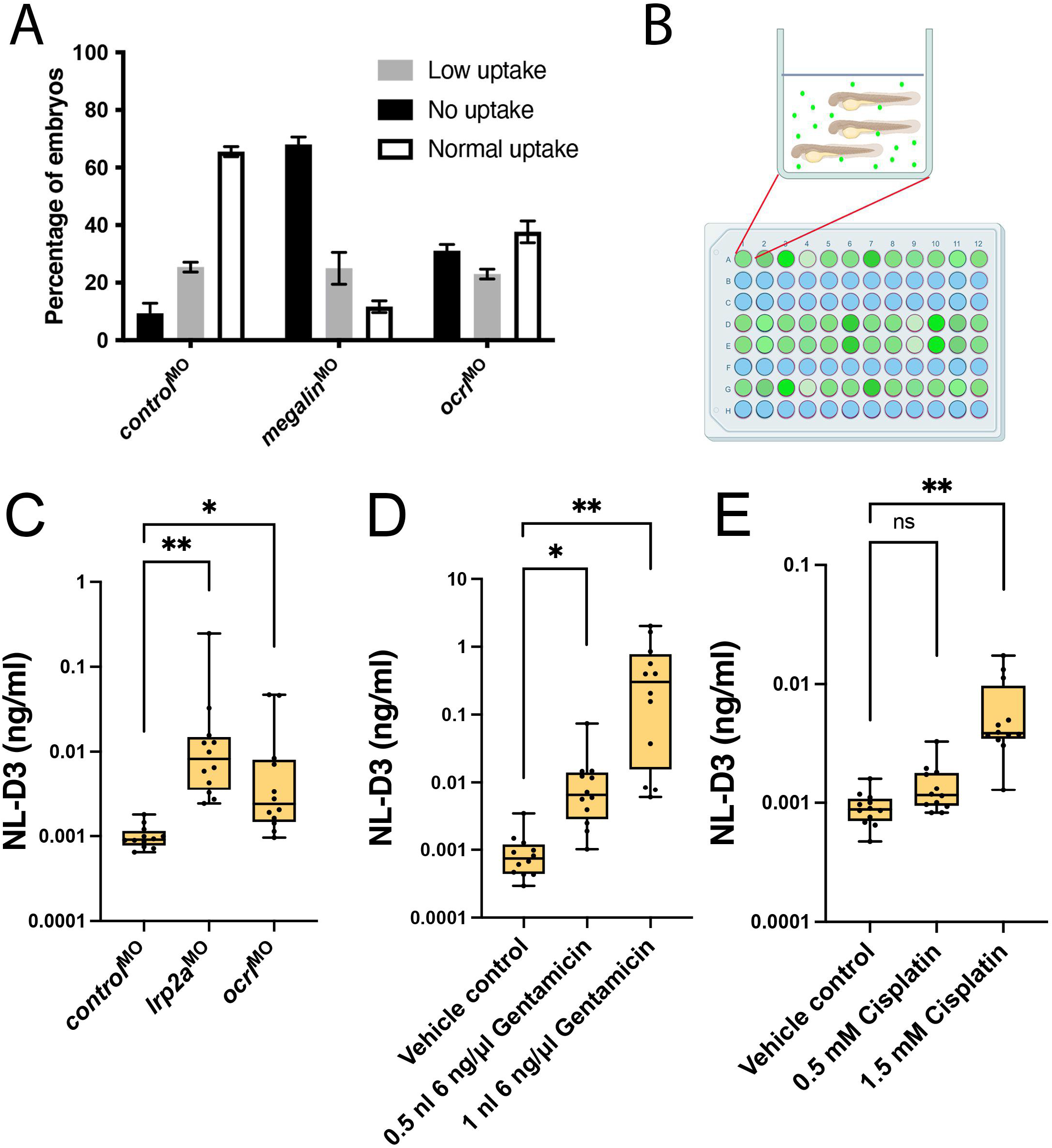
*NL-D3* zebrafish can be used to assay proximal tubule dysfunction. A) Histogram showing the level of uptake of a 10 kDa fluorescent dextran in *control, megalin*, and *ocrl* morphant zebrafish embryos at 5 dpf. B) Schematic showing the experimental setup for the *NL-D3* zebrafish embryos, each 96-well plate was assessed for luminescence on a luminometer. C) Histogram showing the amount of NL-D3 detected in the embryo medium in *control, megalin* and *ocrl* morphants. D) Histogram showing the amount of NL-D3 detected in the embryo medium in vehicle control (DMSO) and two volumes of injected gentamicin. E) Histogram showing the amount of NL-D3 detected in the embryo medium in control and two concentrations of cisplatin.

We further analysed the effects of perturbing the expression of glomerular specific genes to determine whether the *NL-D3* reporter could also be used to measure proteinuria associated with glomerular dysfunction. *NPHS1* and *NPHS2* encode for proteins (nephrin and podocin, respectively) that establish the podocyte slit diaphragm and pathogenic variants in these genes cause nephrotic syndrome, which is characterised by excessive proteinuria. Morpholino-mediated depletion of nephrin and podocin in zebrafish recapitulates phenotypes observed in human patients with disease-causing variants in *NPHS1* and *NPHS2* (Fukuyo et al., 2014). Using identical morpholino oligonucleotides to those used by Fukuyo et al (2014), we observed an increase in NL-D3 in the embryo medium analysed from zebrafish embryos depleted of either *nphs1* (~17-fold increase) or *nphs2* (~11.5-fold increase), (**Figure 3A**). This result demonstrates that the *NL-D3* reporter functions as a dual reporter of both glomerular and proximal tubular dysfunction (**Figure 3B**).

**Figure 3:**
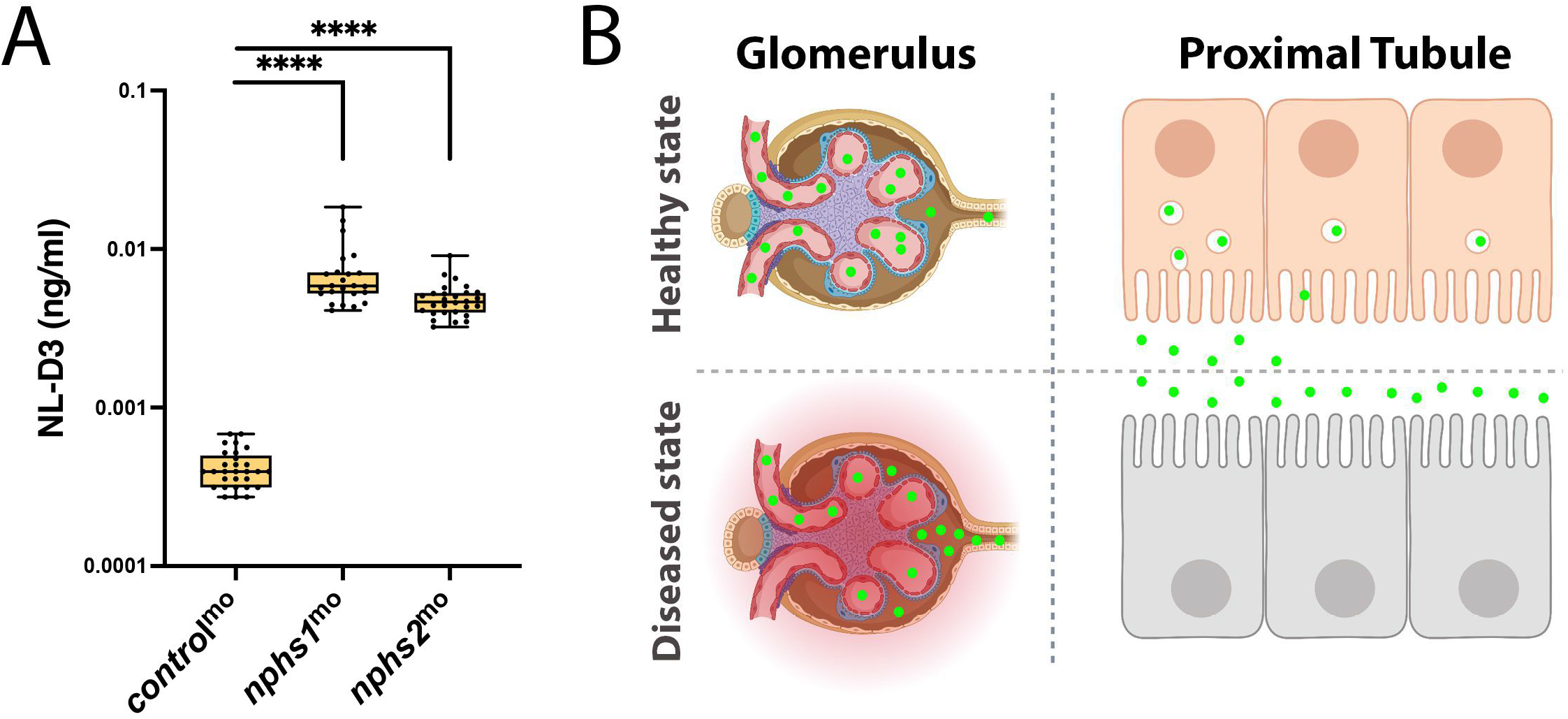
*NL-D3* zebrafish can also be used to study glomerulopathies. A) Histogram showing the amount of NL-D3 detected in the embryo medium in *control, nphs1*, and *nphs2* morphant embryos. B) Schematic (created on Biorender) highlighting the barrier function of the healthy state glomerular filter to NL-D3 and also its re-uptake in healthy state proximal tubules (top panels). The lowered barrier function in the glomerulus and the reduced endocytosis activity in the proximal tubules is displayed in the bottom panels, to schematically represent the dysfunction in these two tissues in a diseased state.

### Expression analysis of type IV collagen genes in zebrafish

The potential for the *NL-D3* reporter line to measure glomerular dysfunction is of particular interest given the plethora of kidney diseases associated with glomerular pathology. One such condition is Alport syndrome, which is caused by variants in *COL4A3/COL4A4/COL4A5* genes in humans. These genes encode for monomers that trimerise to assemble the collagen a3a4a5(IV) network (**Figure 4A**), and which is localized to the mature glomerular basement membrane (GBM). The a3a4a5(IV) network is postulated to provide additional strength to the GBM, allowing it to counteract the high intraglomerular forces generated by capillary wall stress (Gross et al., 2003; Meehan et al., 2009; Savige, 2014). Variants in *COL4A3, COL4A4* or *COL4A5* lead to reduced or absent collagen a3a4a5(IV) trimer in the GBM, affecting its long-term function. Patients with Alport syndrome initially present with microhaematuria, followed by proteinuria and a progressive decline in kidney function leading to kidney failure.

**Figure 4:**
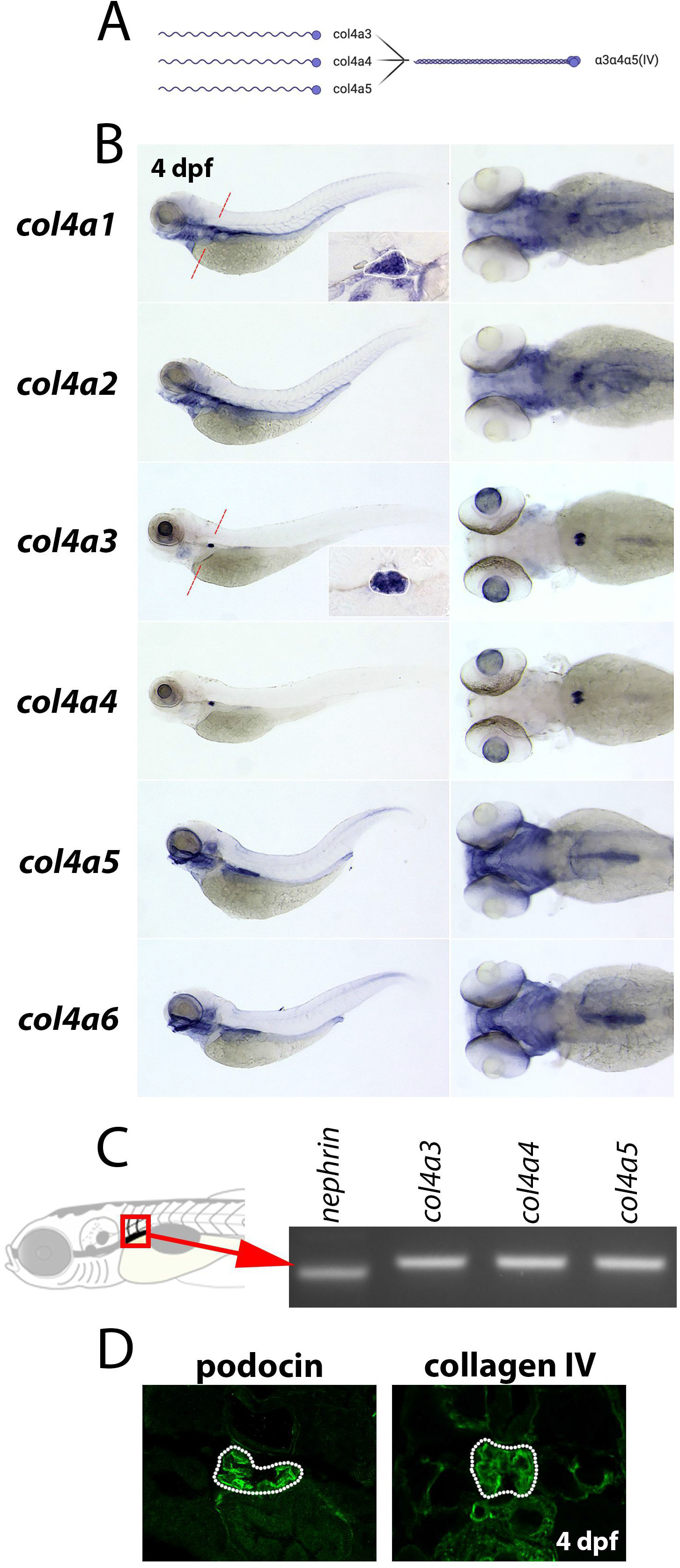
Type IV collagens are expressed in the zebrafish glomerulus. A) Schematic (created with Biorender) showing the three type IV collagen alpha chains expressed in the glomerular basement membrane (α3, α4, and α5) on the left, and their interaction to form the α4α4α5(IV) heterotrimer on the right. B) In situ hybridisations showing the expression patterns of all six type IV collagen chains in zebrafish at 4 dpf. Panels on the left are lateral views of the whole embryo (anterior left). Panels on the right are dorsal views of the head and anterior trunk region (anterior to the left). Inlet panels in the *col4a1* and *col4a3* whole embryo profiles show the glomerulus in transverse section through the point in the embryo highlighted in a red dashed line on the whole embryo image. C) PCR amplicons of *nephrin, col4a3, col4a4* and *col4a5* generated from dissected tissue taken from the glomerular region of the embryo (as shown in the schematic to the left). D) Panels show transverse sections through the glomerulus of 4 dpf zebrafish embryos immunostained for podocin (left panel) and pan-collagen IV (right panel).

The expression profile of type IV collagen chains in the zebrafish glomerulus has not previously been characterised. Zebrafish have six type IV collagen genes, *col4a1-a6*, which are orthologs of the six type IV collagen genes found in humans (*COL4A1-A6*). We performed *in situ* hybridisation to detect the spatial organisation of transcripts for *col4a1-a6* in 4 dpf zebrafish embryos, a time-point when the glomerulus is fully developed (Zhu et al., 2016) (**Figure 4B**). Expression profiles of *col4a1* and *col4a2* were identical, which is expected given type IV collagen gene expression is regulated via putative bidirectional promoters for *col4a1/col4a2, col4a3/col4a4*, and *col4a5/col4a6* (Pöschl et al., 1988; Soininen et al., 1988). *col4a1* and *col4a2* transcripts were detected in the blood vasculature, most notably in the aortic arches, central arteries and primordial hindbrain channels in the head region (**Figure 4B**). Glomerular localisation for *col4a1* was clearly observed in transverse sectioned embryos (**Figure 4B**, inlet). In these sections, *col4a1* also labelled numerous trunk epithelial structures (gut endoderm, pronephric tubules, dorsal aorta, and posterior cardinal vein). When compared to *col4a1* and *col4a2, col4a3* and *col4a4* expression profiles were more restricted. High levels of expression for *col4a3* and *col4a4* were observed in the lens capsule and the glomerulus, as well as weak detection in the branchial arches (**Figure 4B**). Expression of *col4a5* and *col4a6* showed strong localisation for transcripts in the craniofacial region, branchial arches, otic vesicle, developing swim bladder, and posterior gut (**Figure 4B**). Using *in situ* hybridisation we were unable to detect expression of *col4a5* in the glomerulus, despite using different probes. However, PCR analysis on the dissected glomerular region identified *col4a5* transcripts (**Figure 4C**). To assay for the presence of type IV collagen protein in the glomerulus, we performed immunofluorescence with a pan-collagen IV antibody (which detects all six type IV collagen chains) on cryo-sectioned 4 dpf larvae (**Figure 4D**). Localisation of type IV collagen was evident in the glomerulus, as well as in surrounding epithelial tissues.

### *Generation of an Alport syndrome model in the* NL-D3 *zebrafish reporter*

To assay if proteinuria occurs after depleting *col4a3*, *col4a4* or *col4a5* in zebrafish, we adopted a CRISPR-Cas9 approach (Wu et al., 2018) that yields knockout-like phenotypes in F_0_ embryos (termed crispants). This approach robustly induced mutagenesis in *col4a3, col4a4* and *col4a5* when analysed by TIDE analysis (**Figure S3**) and RT-PCR analysis showed transcripts for either *col4a3, col4a4* and *col4a5* were reduced when each gene was targeted using gRNAs (**Figure 5A**). Ultrastructural analysis of the glomerular filtration barrier by transmission electron microscopy identified significant defects in the glomerular capillary wall of the collagen IV crispants (**Figure 5B**). Knockdown of these collagen IV genes reduced the number of podocyte foot processes, created a thicker GBM and precluded normal fenestrated endothelial cell formation (**Figure 5B**). Quantification of average GBM width showed *col4a3* and *col4a4* crispants had thicker GBMs within 95% confidence limits, and *col4a5* crispants also had a thicker GBM within 92% confidence limits (**Figure 5C**). Similarly, measurement of average foot process length showed that depletion of each collagen IV gene increased the width of podocyte foot processes, indicating effacement (**Figure 5D**). These glomerular ultrastructure changes phenocopy those observed in human patients with Alport syndrome and knockout mouse models of Alport syndrome (Cosgrove et al., 1996; Miner and Sanes, 1996).

**Figure 5:**
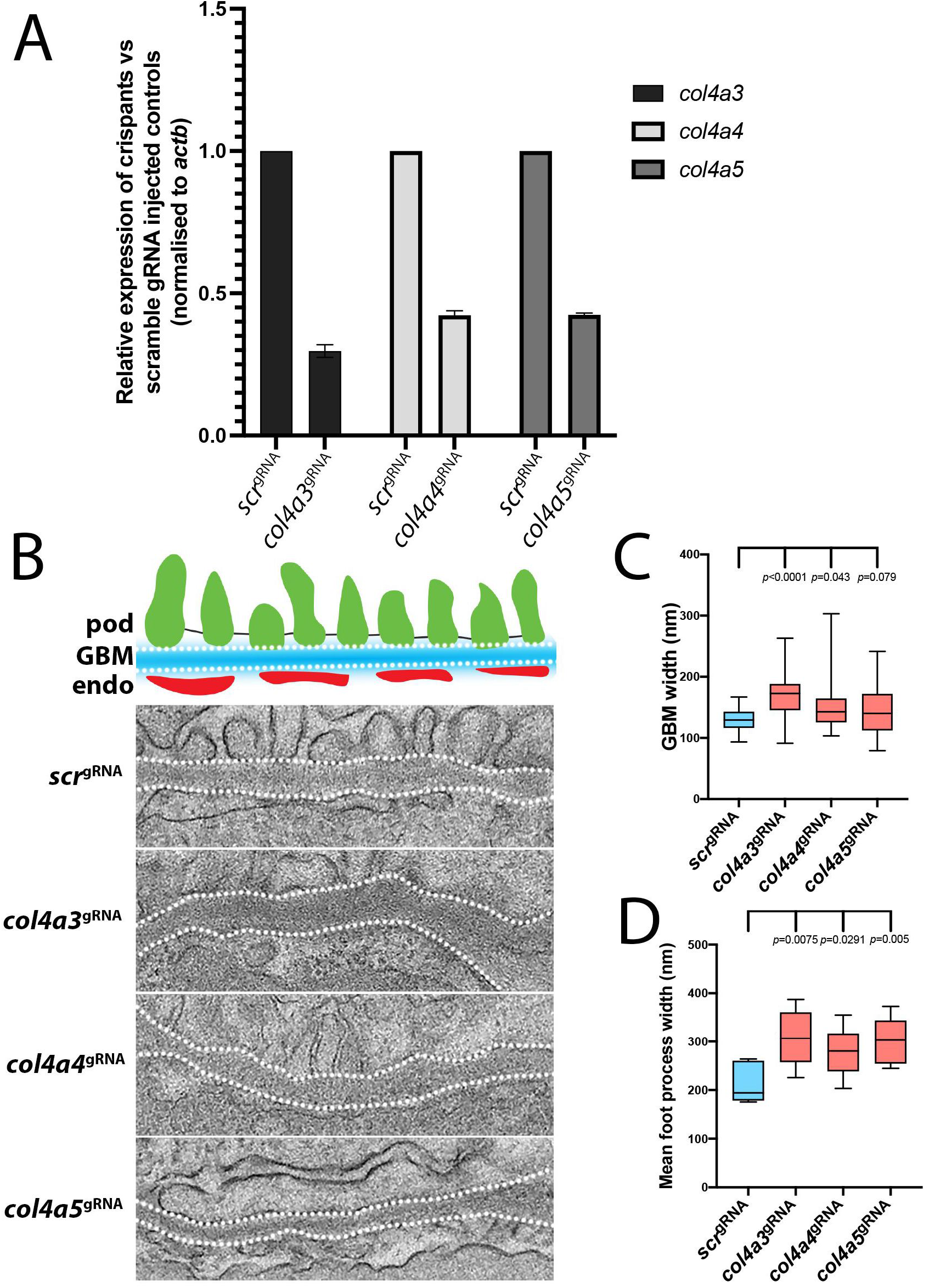
Depletion of *col4a3, col4a4* or *col4a5* induces Alport syndrome phenotypes in zebrafish. A) Histogram showing RT-PCR analysis of *col4a3, col4a4* and *col4a5* in wild-type versus respective crispant embryos. B) Top panel shows a schematic representation of a cross-section through a glomerular filtration barrier (Abbreviations: pod, podocyte foot process; GBM, glomerular basement membrane; endo, fenestrated endothelium). Below panels show representative images of the glomerular filtration barrier in control *scr*, *col4a3, col4a4* and *col4a5* crispants. The white-dotted lines indicate the GBM, with the podocyte processes above and endothelial cells below. C) Histogram showing the average GBM width in control *scr*, *col4a3, col4a4* and *col4a5* crispants. D) Histogram showing the average foot process width in control *scr*, *col4a3, col4a4* and *col4a5* crispants.

To determine if depletion of *col4a3*, *col4a4* or *col4a5* caused proteinuria, we injected gRNAs targeting these genes into our transgenic *NL-D3* reporter line. Injected embryos were grown to 4 dpf and then placed in shoals of 3 embryos in individual wells of a 96-well plate. The embryo medium was replaced, and the embryos were left for 24 hours to accumulate NL-D3 for quantitative analysis of luminescence. This analysis showed that knockdown of each of the three Alport genes in zebrafish crispants induced proteinuria (~12-fold for *col4a3* crispants, ~7.4-fold for *col4a4* crispants, and ~6-fold for *col4a5* crispants) when compared to *control*^gRNA^-injected embryos (**Figure 6A**). In conclusion, knockdown of either *col4a3, col4a4* or *col4a5* caused ultrastructural defects in the glomerular filtration barrier and induced proteinuria, recapitulating the phenotypes of pathogenic variants in Alport genes in both mouse knockout models and human patients.

**Figure 6:**
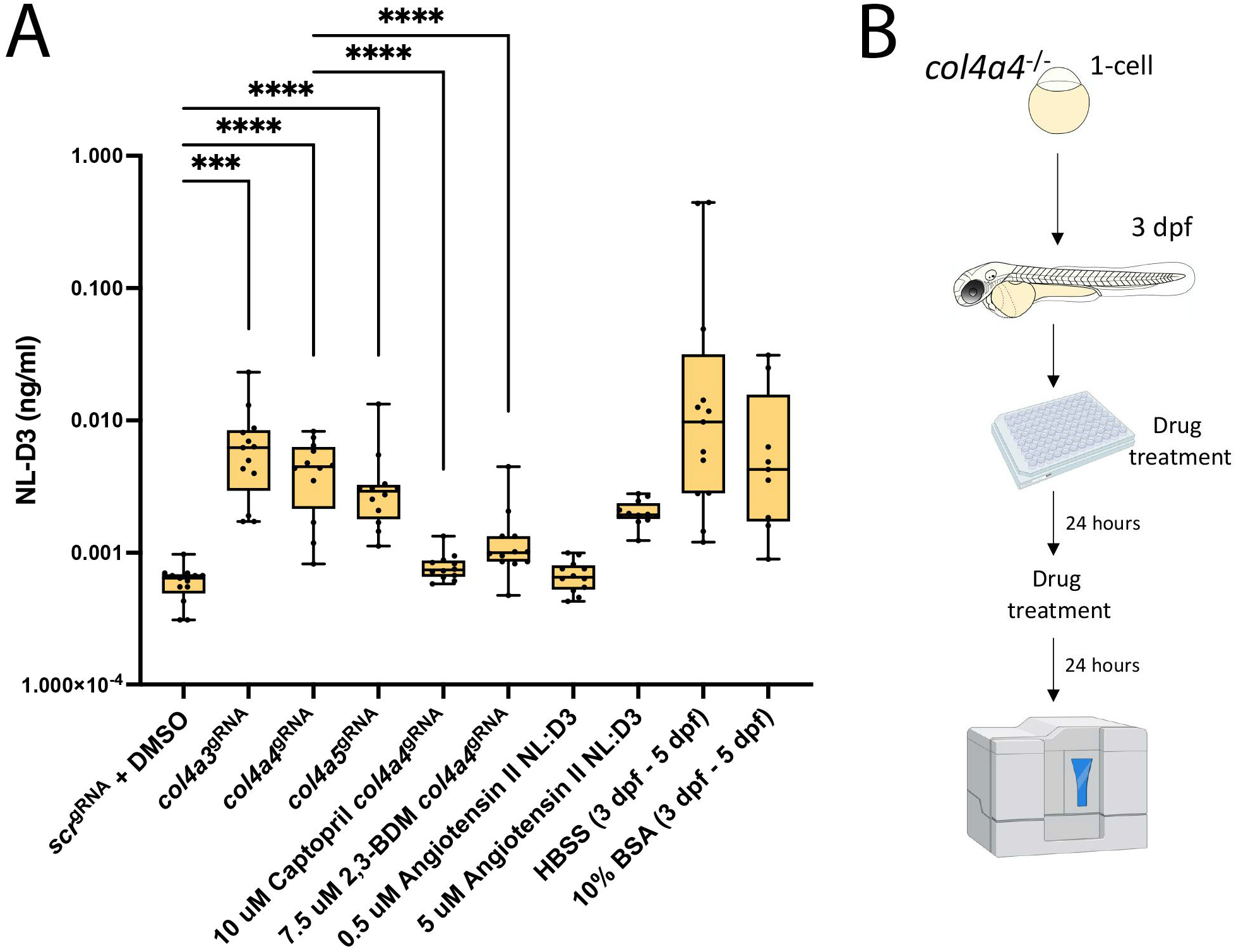
Alport zebrafish display proteinuria which can be alleviated by reducing intraglomerular force. A) Histogram showing amounts of NL-D3 produced into embryo medium in control *scr*, *col4a3, col4a4* and *col4a5* crispants. The effect of the angiotensin converting enzyme inhibitor (ACEi) captopril and 2,3-Butanedione monoxime (2,3-BDM) on proteinuria in *col4a4* crispants is also shown. Angiotensin II, HBSS, and 10% BSA all increased proteinuria in *NL-D3* zebrafish embryos. B) Schematic showing the experimental pipeline for assaying the effects of chemicals in zebrafish *NL-D3* embryos.

### Pharmacological amelioration of proteinuria in zebrafish Alport crispants

Given the zebrafish *NL-D3* transgenic reporter line can be used for assaying glomerular dysfunction where proteinuria is a key clinical feature, such as in Alport syndrome, we next attempted to rescue glomerular filtration using chemical approaches to determine the potential for this system for compound screening. In *col4a4* crispants from 3 dpf to 5 dpf, we used captopril to reduce the systemic blood pressure, and we assayed for proteinuria from 4 dpf to 5 dpf (**Figure 6B**). Captopril is an angiotensin-converting enzyme inhibitor (ACEi) and has previously been used in zebrafish to lower blood pressure (Rider et al., 2015). We found that captopril treatment had no effect on un-injected zebrafish embryos (data not shown) but almost entirely prevented proteinuria in *col4a4* crispants (**Figure 6A**). We also used 2,3-butanedione 2-monoxime (2,3-BDM) to slow the heart rate in zebrafish embryos from 3 dpf to 5 dpf. We hypothesised that a lower heart rate would lead to reduced glomerular perfusion, thereby reducing glomerular filtration. In wild-type zebrafish embryos, a dose response analysis showed 2,3-BDM at a concentration of 7.5 μM was sufficient to significantly lower heart rate (**Figure S4**). Incubation of *col4a4* crispants with this concentration of 2,3-BDM lowered proteinuria in these animals to a level that was still ~2.2-fold higher than controls but ~3.3-fold lower than untreated *col4a4* crispants (**Figure 6A**).

Using the *NL-D3* reporter line, we were also able to detect the effects of increasing systemic blood pressure on zebrafish embryos. Angiotensin II is a peptide hormone that increases blood pressure by inducing vasoconstriction. We found that 0.5 μM angiotensin II was unable to induce proteinuria in *NL-D3* embryos, however 5 μM angiotensin II resulted in a ~3.4-fold increase in proteinuria. Similarly, injection of Hank’s Balanced Salt Solution (HBSS) or 10% Bovine Serum Albumin (BSA) caused a dramatic increase in proteinuria in the *NL-D3* reporter embryos. HBSS is a saline solution that upon injection causes water influx into the blood vasculature. Thus, we hypothesise that the injection of HBSS creates an acute mechanical loading in the glomerulus that we predicted would cause proteinuria. In support of this we observed glomerular swelling after HBSS injection in transgenic *nphs2:egfp* embryos that have podocytes labelled with eGFP (data not shown). We detected a ~128-fold increase in luminescence in HBSS-injected embryos (**Figure 6A**). Human albumin solution is given in the clinical setting to expand the circulation and thus we injected a high concentration BSA solution (10%) into zebrafish embryos to increase glomerular perfusion. We again observed an increase in proteinuria, which was ~15-fold higher than non-treated controls (**Figure 6A**).

Overall, these data confirm the ability of our *NL-D3* reporter line to detect proteinuria associated with changes in systemic blood pressure. They also highlight the potential of our new tool to be used to develop or screen for drugs that affect glomerular function in normal or diseased zebrafish embryos.

## Discussion

Here, we describe a new high-throughput tool for the quantification of proteinuria in the zebrafish model. We find that the *NL-D3* transgenic reporter can detect changes in both glomerular and proximal tubular function. Our investigations into the impact of chemical agents on wild-type and genetically modified kidneys show that significant changes in proteinuria can be detected. Furthermore, dose-dependent responses were observed in gentamicin, cisplatin and angiotensin II treatments. Given our *NL-D3* transgenic reporter is a simple system, it is attractive for future mutation and drug compound screening. The use of nanoluciferase also ensures this reporter is easy to adopt by research groups as luminometers are commonly found in most laboratories. To aid future users of the line, we generated a standard curve to enable conversion of relative luminescence units (RLUs) to NL-D3 amount in ng/ml. Conversion of RLUs to NL-D3 amount is not fundamental to infer changes in proteinuria in the reporter, but our empirical standard curve data is provided in Supplementary File 1, with which users can interpolate their RLU readings in GraphPad Prism and generate NL-D3 ng/ml readings.

We robustly assessed the *NL-D3* reporter to ascertain any limitations and considerations with the line. We have found that obtaining reliable data is dependent on maintaining the same parameters and equipment when collecting the luminescence readings. Given the reporter is a transgenic system, there is likely to be animal-to-animal and generation-to-generation variability in expression of the transgene. To reduce the impact of transgene expression variation, we isolated embryos that had high levels of mCherry fluorescence in the lens before analysis or drug treatments began (at 3 dpf). We also found that placing three embryos per well in a 96-well plate, instead of one embryo, reduced variability. After taking these measures the variability in wild-type embryos was negligible (deviation less than 10% from the mean). We did detect greater variability in the levels of proteinuria in embryos treated with chemicals or whose gene products were knocked down. This is likely to be a consequence of dose variation of chemical, morpholino or gRNA rather than variation in transgene expression. Even with these variations, we were able to detect statistically significant changes in proteinuria in the experimental samples. The brightness of the nanoluciferase assay system also means that if any embryos die or are accidentally destroyed during the experiment, the embryo medium from these wells cannot be read (they commonly gave luminescence readings of >10,000-fold higher than controls) and can therefore be reliably excluded from the data. To reduce the impact of high readings, we also used opaque 96-well plates to minimise any bleed-through from luminescence in neighbouring wells.

The luminescence scores in our lysate analyses or in wells where embryos had deceased were high compared to untreated control embryos. This highlights the impressive ability of the embryo to retain the reporter internally. We expected the NL-D3 protein to readily pass through the glomerular filter and be required to be reabsorbed in the proximal tubule via megalin-mediated endocytosis. In support of this, impeding the proximal tubule endocytosis re-uptake pathway by morpholino-knockdown of *lrp2a* or *ocrl* caused proteinuria and so confirmed the proximal tubules, in a healthy state, can reabsorb most of the NL-D3 that passes through the glomerulus. However, the glomerulus must also act as a partial barrier to NL-D3 given that we find genetic or chemical targeting of glomerular filtration induces an increase in the presence of NL-D3 in the embryo medium. It is possible that the negative charge of the NL-D3 protein (given its predicted isoelectric point of ~6.29) may impede its transition through the glomerular filter, which similarly has a net negative charge. However, the significance of charge selectivity in glomerulus remains unknown (Brenner et al., 1978; Goldberg et al., 2009; Khalil et al., 2019; van den Hoven et al., 2008). The lower-than-expected filtration of NL-D3 through the glomerulus would mimic the glomerular barrier function to serum albumin, which was thought to be small enough to pass through the filter, but has a low sieving coefficient (0.00062) (Tojo and Endou, 1992). Slit diaphragm pore size is roughly equivalent to the size of serum albumin (Gekle, 2005; Wartiovaara et al., 2004), yet it is unable to pass through the filter readily. Recent evidence suggests that the podocyte glycocalyx and the extent of GBM compression are important for the size selectivity of the glomerular filter (Butt et al., 2020; Lawrence et al., 2017). Thus, we speculate that these barriers prevent passage of NL-D3 via simple diffusion through the glomerular filter in the healthy state. However, when podocyte effacement occurs (as we see in our TEM images of Alport crispant zebrafish) the GBM compression and slit diaphragm are lost and NL-D3 is more readily able to pass through the filter, saturating the reabsorption pathways in the tubules, and leading to the proteinuria we detect.

In our studies we developed a new model of Alport syndrome that phenocopies many of the disease features found in human patients. The comparative conservation of morphological and molecular features in the human and zebrafish glomerulus (Ichimura et al., 2012; Wingert et al., 2007) means our zebrafish proteinuria reporter is an ideal model with which to investigate Alport syndrome. We have shown that *col4a3* and *col4a4* are expressed in the zebrafish glomerulus by *in situ* hybridisation and PCR analysis. We were unable to detect *col4a5* by *in situ* hybridisation, which is likely to be due to the technical difficulties (such as probe penetration). Dissection of pure glomerular tissue was not possible due to the adhesiveness of this organelle to its surrounding tissues. However, dissection of the region around the glomerulus was successful and *col4a5* transcripts were detected. We cannot absolutely rule out contamination from surrounding tissue accounts for some or all this transcript expression. However, we believe it is very likely that *col4a5* is expressed in the zebrafish glomerulus. The stoichiometry of the α3α4α5(IV) heterotrimer is well characterised in humans (Kobayashi and Uchiyama, 2003; Omachi et al., 2018; Pedchenko et al., 2021) and the zebrafish α3, α4 and α5 chains share sequence homology with the GXY collagenous domain present in the orthologous human chains. Thus, it seems unlikely that the zebrafish forms a heterotrimer combination that is dissimilar to the human type IV collagens (either α1α1α2(IV), α3α4α5(IV), or α5α5α6(IV)). Further to this, we found that depletion of either *col4a3, col4a4* or *col4a5* caused glomerular ultrastructural changes and proteinuria, demonstrating that loss of either chain results in disease features associated with Alport syndrome. For this study, we used a combinatorial gRNA approach to introduce indels in F_0_ crispant embryos (Wu et al., 2018), which is not suited to establishing stable lines due to each gene being exposed to 4 gRNAs, making genotyping exhaustive and increasing the chance of off-target effects. However, we did grow these fish to breeding age and did not detect any increase in mortality rates over un-injected controls (**Figure S5**). Thus, in the future a stable set of Alport zebrafish lines will be viable to develop. Such a stable line could be used for a drug screen to discover chemicals that ameliorate proteinuria associated with Alport syndrome. We have shown that captopril and 2,3-BDM are able to lower proteinuria back to healthy levels, demonstrating the potential of this new tool for screening purposes.

In summary, our data shows the development of a nanoluciferase based reporter for proteinuria in the zebrafish system. Given the advantages of this system in terms of experimental usability and the feasibility of high-throughput screens, we anticipate this new proteinuria reporter will be a valuable addition to the toolkit of kidney researchers.

## Author contributions

M.L, A.J-C and E.L. designed and generated the NL-D3 transgenic reporter. R.L. and R.W.N. generated the Alport zebrafish crispants (with the expert help of Dr. Antony Adamson of the University of Manchester GeU). B.D. prepared the Alport zebrafish embryos for EM analysis and A.M. carried out the E.M. analysis. R.W.N., E.L., R.L. and M.L. wrote the manuscript.

## Acknowledgements

This work was supported by a Wellcome Trust Senior Fellowship awarded (202860/Z/16/Z) to R.L., supporting R.W.N., and B. D., and Lowe Syndrome Trust grants awarded (ML/MU/2012 and ML/MU/2016/2017) to M.L, supporting A.J-C and E.L., respectively. The authors thank the staff in the BSF unit for help with zebrafish maintenance. The Bioimaging Facility microscopes used in this study were purchased with grants from BBSRC, Wellcome and the University of Manchester Strategic Fund. Special thanks goes to Peter March, Roger Meadows *and* Steven Marsden for their help with the microscopy. The authors thank the staff in the EM Core Facility in the Faculty of Biology, Medicine and Health for their assistance, and the Wellcome Trust for equipment grant support to the EM Core Facility

---

**Figure S1:**
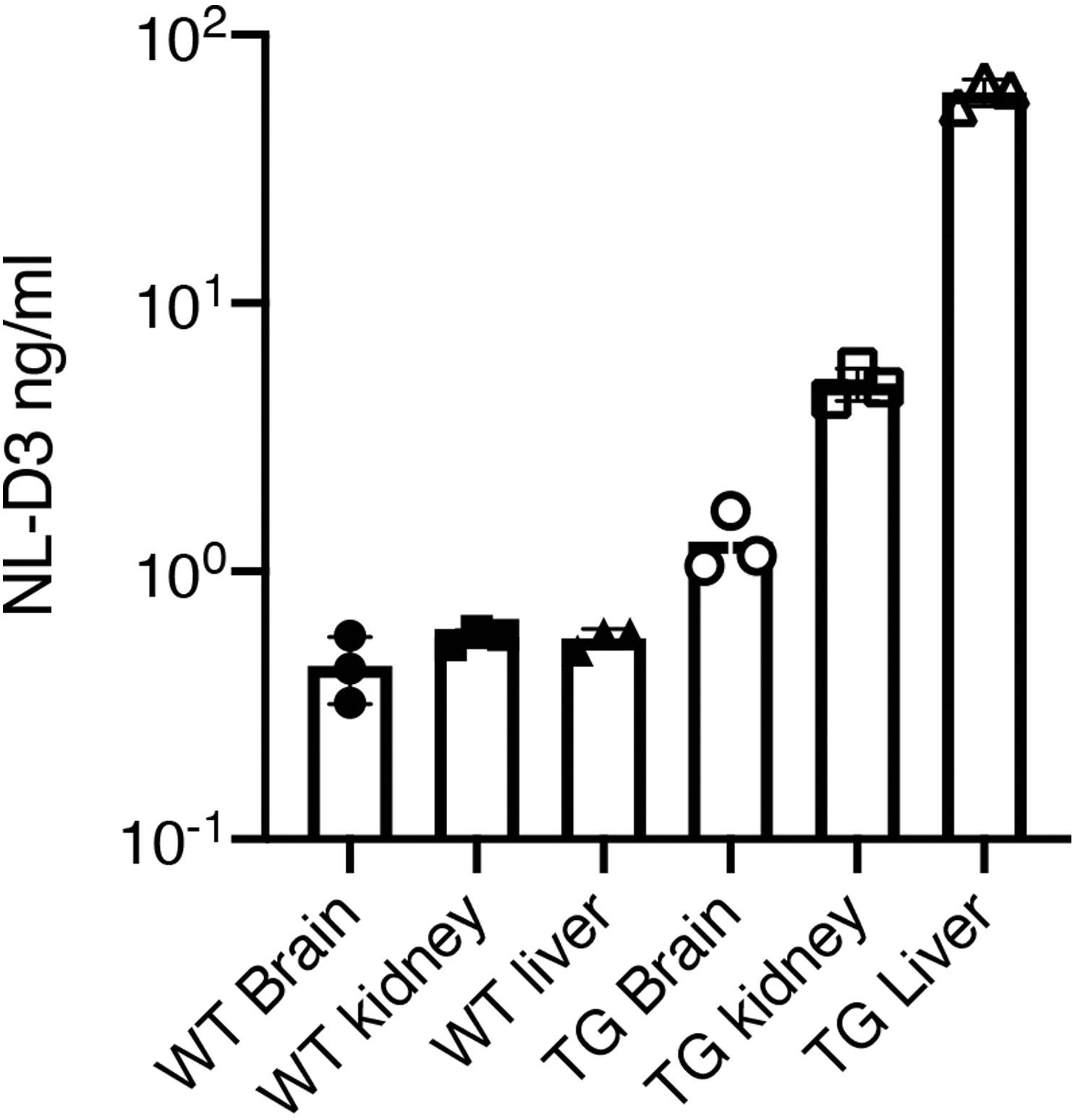
Comparison of luminescence in wild-type versus *NL-D3* adult tissue. Histogram showing the relative luminescence intensities of brain, kidney and liver tissue extracted from wild-type (WT) and *NL-D3* (TG) adult zebrafish.

**Figure S2:**
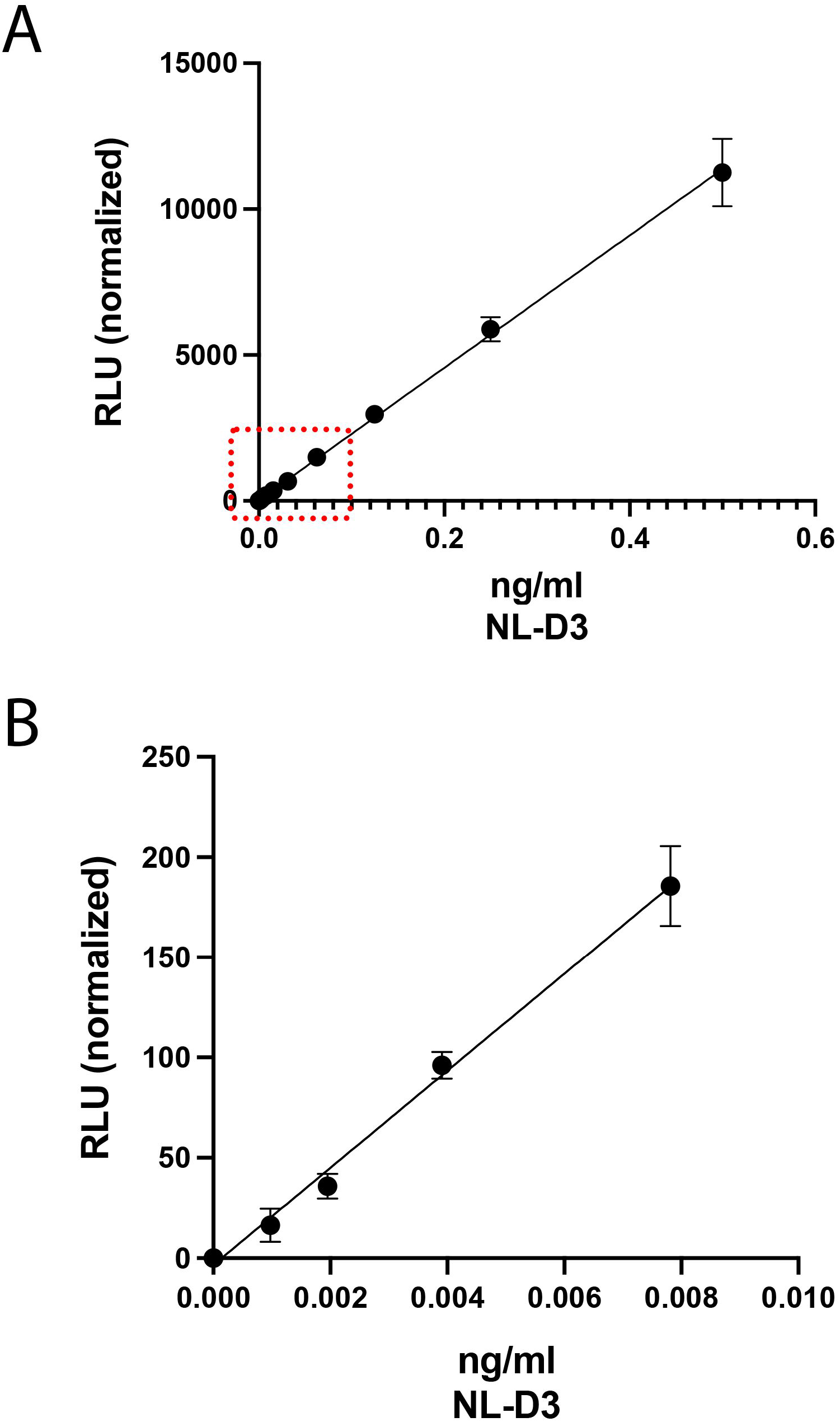
Standard curve data. A) Graph showing standard curve curated from data collected analysing 0.5 ng/ul to 0.0005 ng/ul. B) Close up visualisation of the graph in (A) in the region of the red-dotted box to highlight the lower concentrations of NL-D3 where most data points were collected.

**Figure S3:**
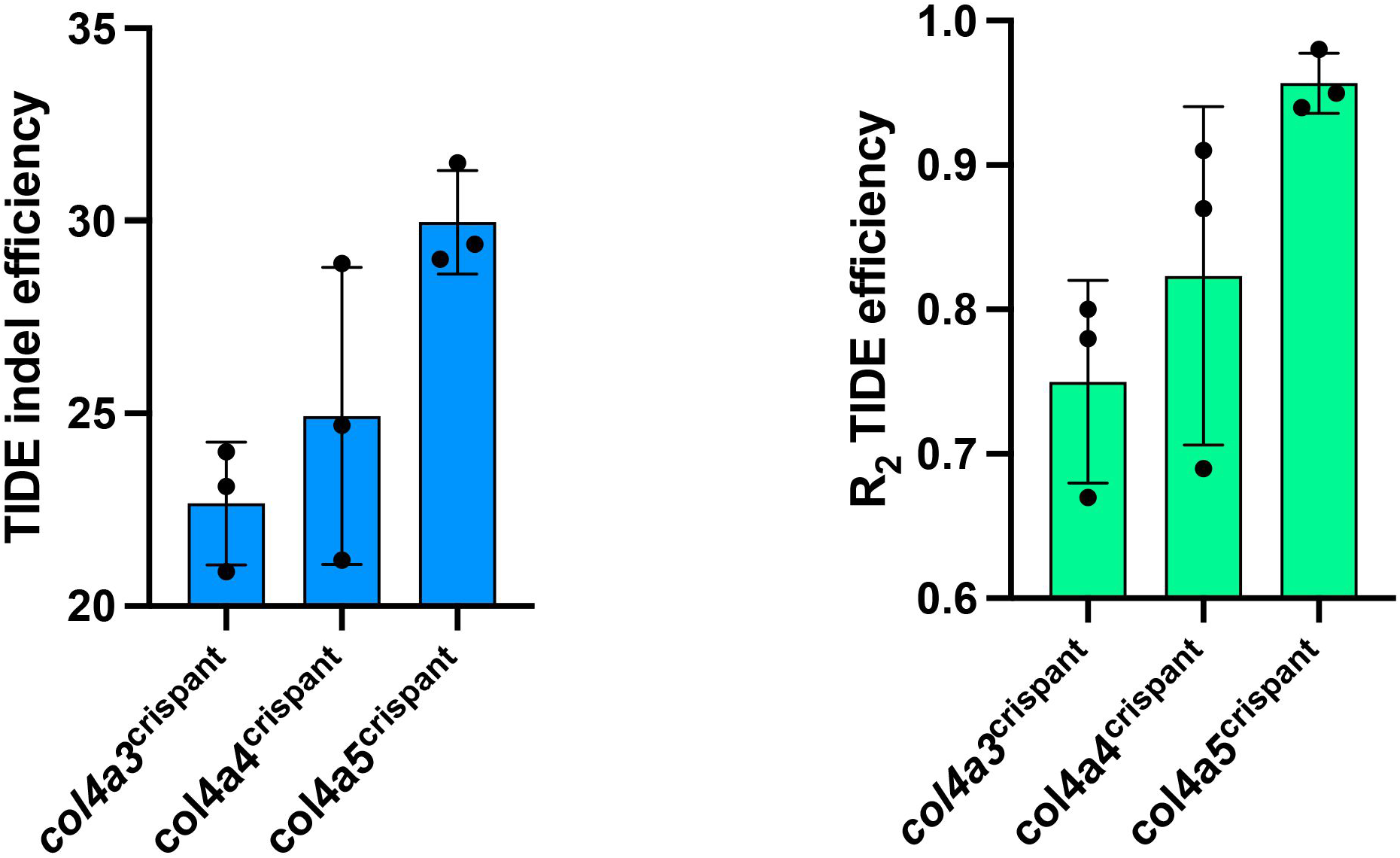
TIDE analysis shows mutagenesis at gRNA site 1 in the Alport zebrafish crispants. Histogram on the left shows the TIDE indel efficiency (calculated with the algorithm described on https://tide.nki.nl/) for three biological replicates induced by gRNA 1 for the col IV crispant zebrafish embryos. Histogram on the right shows the R2 values for the corresponding indel efficiencies.

**Figure S4:**
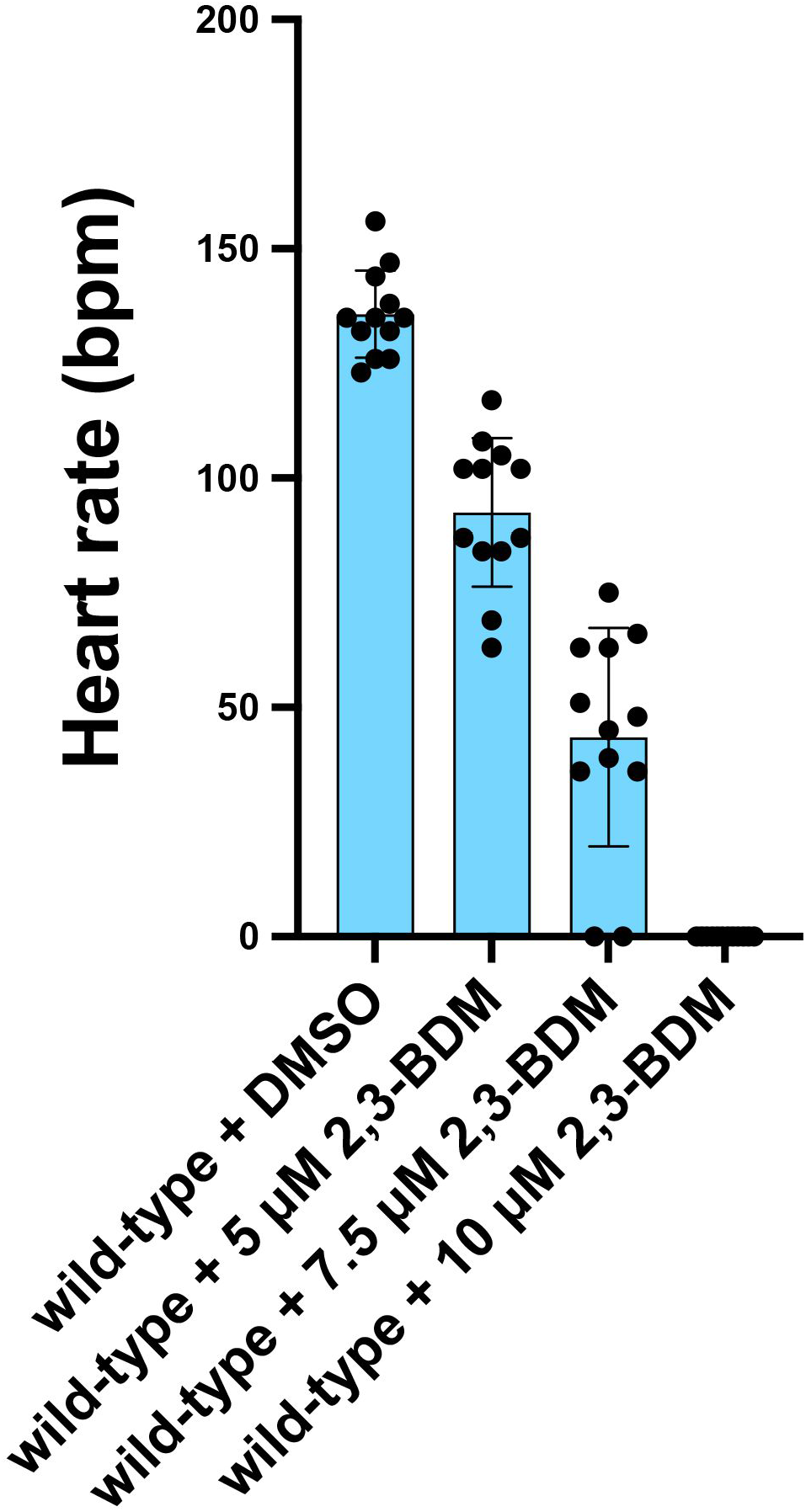
Increasing 2,3-BDM dose results in increasing effect on heart rate in zebrafish embryos. Histogram shows heart rate (bpm) of wild-type embryos treated for 1 hour with vehicle (DMSO) or 2,3-BDM of varying concentrations highlighted.

**Figure S5:**
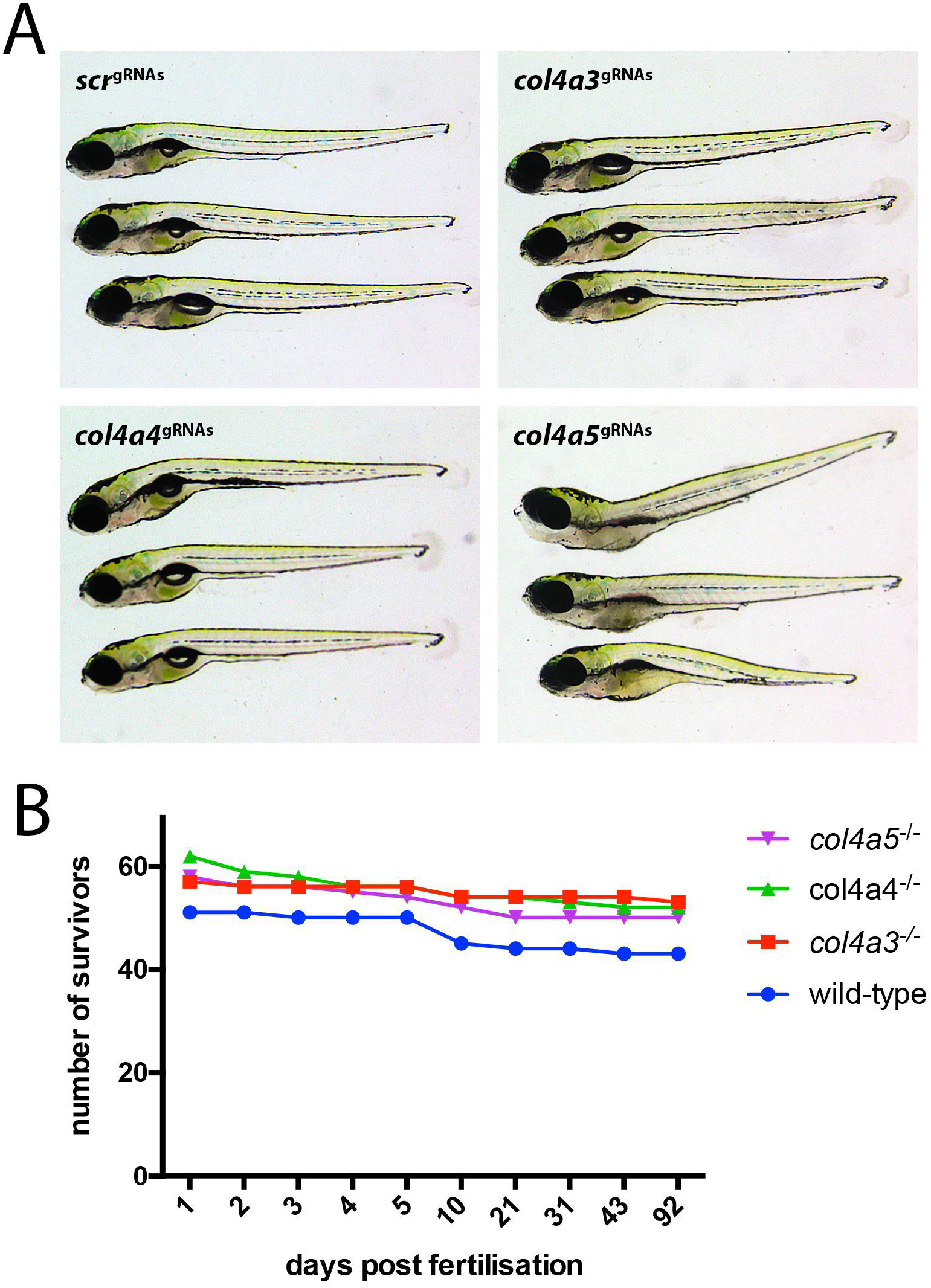
Alport crispants survive to adulthood. A) Panels show three zebrafish Alport crispant embryos in lateral view (anterior to left) injected with the displayed gRNAs. B) Graphical representation of survival rate of Alport zebrafish crispants to breeding age.

